# Computational simulation reveals the critical role of spike-timing-dependent plasticity in synchrony

**DOI:** 10.64898/2026.08.27.747601

**Authors:** Yunqi Huang, Milad Lankarany, Gabriele D’Eleuterio

## Abstract

Emerging research on desynchronization reveals strong relations to memory formation and recollection with much of the evidence originating from studies of the hippocampus, a brain region central to memory processing. However, growing evidence emphasizing the role of desynchronization is accompanied by challenges, most notably the synchronization/desynchronization conundrum. The conundrum arises from an apparent contradiction between learning and Shannon’s information theory: spike-timing-dependent plasticity (STDP) acquires information via synchronized populational activity whereas information theory suggests that information channel is encoded through higher variability. Both frameworks are well-founded. Learning is widely accepted as the mathematical abstraction of STDP while Shannon’s information theory showcases its power ubiquitously across fields from communication systems to neuroscience. Here, we propose that learning and Shannon’s information theory are not necessarily opposing principles. Rather, learning plays a critical role in the process of synchronization and desynchronization. To justify our claim, we conduct computational simulations based on a Hodgkin-Huxley neuronal model coupled with NMDA, AMPA, and GABA synapses with inputs of biologically realistic spiking data. Our results show that enabling STDP significantly influences synchronization and desynchronization dynamics. Specifically, we find that STDP and desynchronization form a regulatory loop, in which STDP regulates the level of synchrony, and synchrony regulates subsequent learning strength. Further study reveals STDP is capable of switching neurons from synchronization to desynchronization. This framework mitigates the synchronization/desynchronization conundrum by clarifying the relationship between desynchronization and learning, and offers a new perspective on how synchrony contributes to memory formation and recollection.

## 1 Introduction

Episodic memory involves encoding of events and retrieval of events and their contextual details [1]. Its formation and recollection are accompanied by brain oscillations [2]. Oscillations of the theta band (3 *−* 10 Hz) and gamma band (40 *−* 80 Hz) are the embodiment of synchronization within a neuronal population, where different neurons spike in a coordinated behaviour. Such synchronization has been observed across multiple brain regions, including the neocortex (alpha oscillations) and the hippocampus (theta oscillations) [3, 4]. Both regions are known to be highly involved in cognitive functionality such as working memory and attention to spatial and feature attributes [5, 6].

Earlier studies have primarily explored the role of synchronization in cognition. State-of-the-art research offers growing evidence highlighting the functional significance of desynchronization within neuronal assemblies [7]. Desynchronization has been shown to play a critical role in memory encoding and retrieval. Across studies, the types of memory examined vary from pattern recognition to context-related communication signals [8, 9]. Computational studies seek to create models to reproduce the oscillatory behaviour and, ultimately, to dissect the mechanisms of initiation or disruption of oscillations during memory acquisition and recollection [10].

Despite extensive research supporting the significance of desynchronization, one central issue remains: the synchronization/desynchronization conundrum [2]. The conundrum arises from the contradiction between Hebbian learning [11] — summarized by the statement “neurons that fire together, wire together” — and Shannon’s information theory. Hebbian learning is biologically supported by the experimental studies on long-term potentiation (LTP) and long-term depression (LTD), together forming spike-timing dependent plasticity (STDP) [12]. Experimental studies have discovered long-term potentiation in multiple brain regions, including the hippocampus [13]. For glutamatergic synapses, both LTP and LTD are predicated on the activation of N-methyl-daspartate (NMDA) receptors [14]. In addition, *α*-amino-3-hydroxy-5-methyl-4isoxazolepropionate (AMPA) receptor traffic is also found to influence critically LTP and LTD [15, 16]. LTP and LTD are not exclusive to the excitatory synapse. In fact, *γ*-aminobutyric acid (GABA)-ergic synapse also exhibits both forms of plasticity [17]. The prevalence of LTP and LTD across diverse synapses suggests that STDP is broadly applicable. The synchronous firing of presynaptic neurons and postsynaptic neurons forms a key portal of information acquisition. In contrast, information theory argues that information is conveyed through variability. Probability is the foundation of information entropy. Stochasticity or variability is required to encode information by increasing channel capacity. When applied to STDP, this perspective suggests that synchronization reduces the amount of information, seemingly contradicting the purpose of learning information. This discrepancy between neurophysiological and probabilistic perspective reflects a need to study further the mechanism behind learning.

We propose a novel idea that can reconcile the conflict. Specifically, we posit the existence of a control loop between STDP and synchrony where STDP tunes synchrony and, in turn, synchrony controls the strength of STDP. The proposal is further demonstrated using a computational model composed of Hodgkin-Huxley neurons and simulated NMDA, AMPA and GABA synapse. The model simulates the key neuronal mechanism of the hippocampus and performs STDP on all three types of synapses, allowing us to isolate synaptic changes arising solely from STDP mechanisms.

## 2 Methods

The computational model is designed to demonstrate the proposed mechanism whereby STDP drives synchrony. To this end, we perform a comparative computational simulation using the same network of neurons (Figure 1), with STDP either enabled or disabled.

**Fig. 1.**
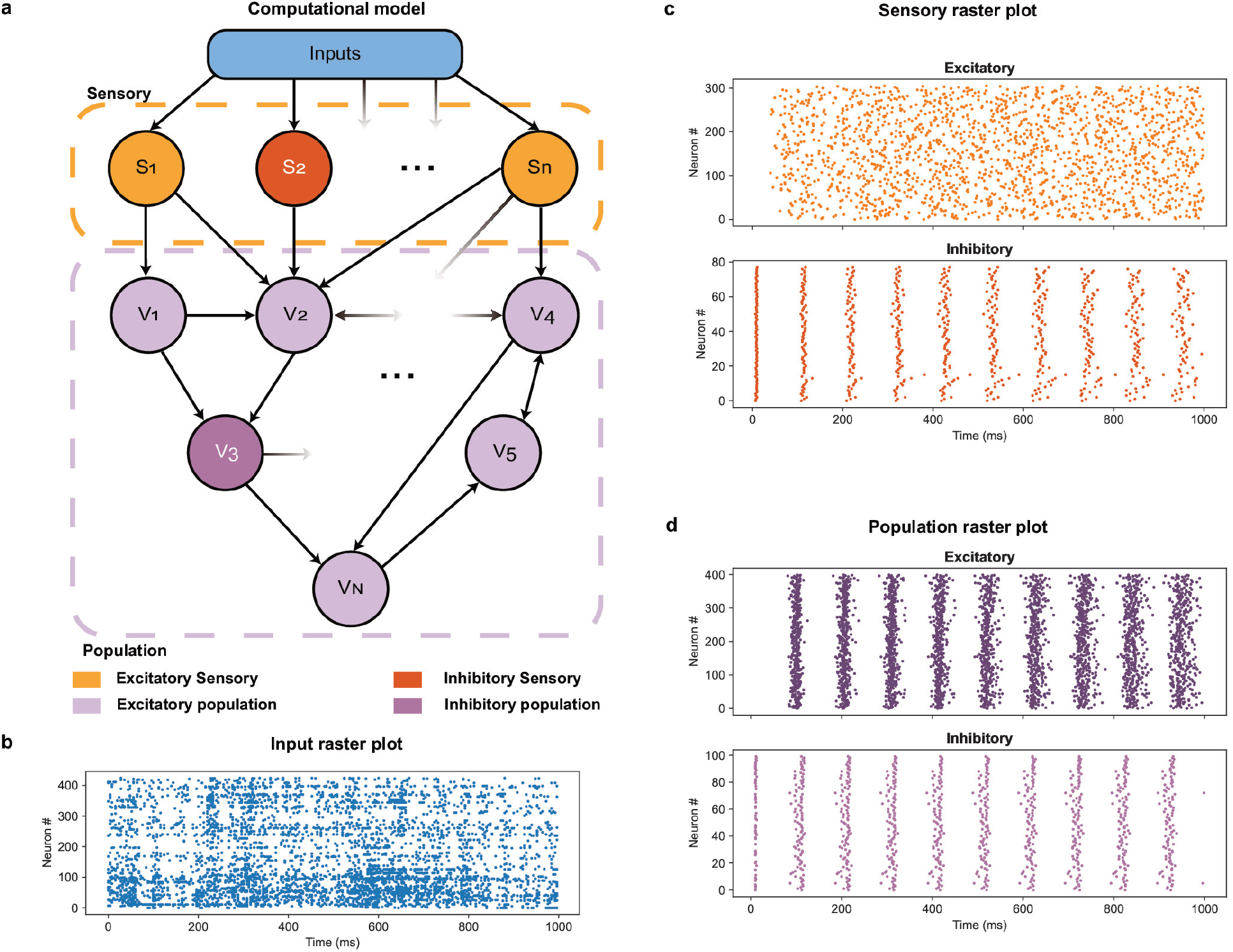
Illustration of computational simulation: **a**. Model schematic. The model is composed of three layers: input, sensory and population layer. **b-d** Raster plot of spiking neurons in each layer with the separation of excitatory and inhibitory for sensory and population layer.

### 2.1 Model structure

The model is composed of two different layers: a sensory layer and a population layer. The input is defined according to the input dataset provided. It is expected to include a specific number of neurons and the recording of the spiking events over time. The inputs are then provided to the sensory layer containing the same number of neurons as the input. This layer mainly models the reception of these inputs and further passes these inputs to the population layer, which is composed of 500 neurons. Since the neurons in both sensory and population layers are created artificially, we randomly assign these neurons to 80% excitatory and 20% inhibitory based on rough statistics of visual cortex neurons’ polarity percentage [18].

Synapses between and within these layers are formulated with random assignments but following these rules. The connections between sensory and population layers are unidirectional: the sensory layer receives only input spikes and the population layer from the sensory layer. The intralayer connections only exist for population layer. For connections to population neurons, we group sensory and population neurons of the same polarity together. Each population neuron randomly chooses 10% from each polarity group and assigns them as presynaptic neurons, based on average measurement data [19].

### 2.2 Input dataset

The input of the simulation uses rat neuronal and behavioural dataset measured when performing a visual contrast task [20]. The head-fixed rats are shown with two images with different contrast. They are trained to choose the image with higher contrast by turning the wheel in hand towards it (Figure 2). When turning in the correct direction, the rat is rewarded with currant juice. In the meantime, spike data are recorded by electrodes in multiple brain area. Spikes are then sorted to each neuron and fed into sensory inputs.

**Fig. 2.**
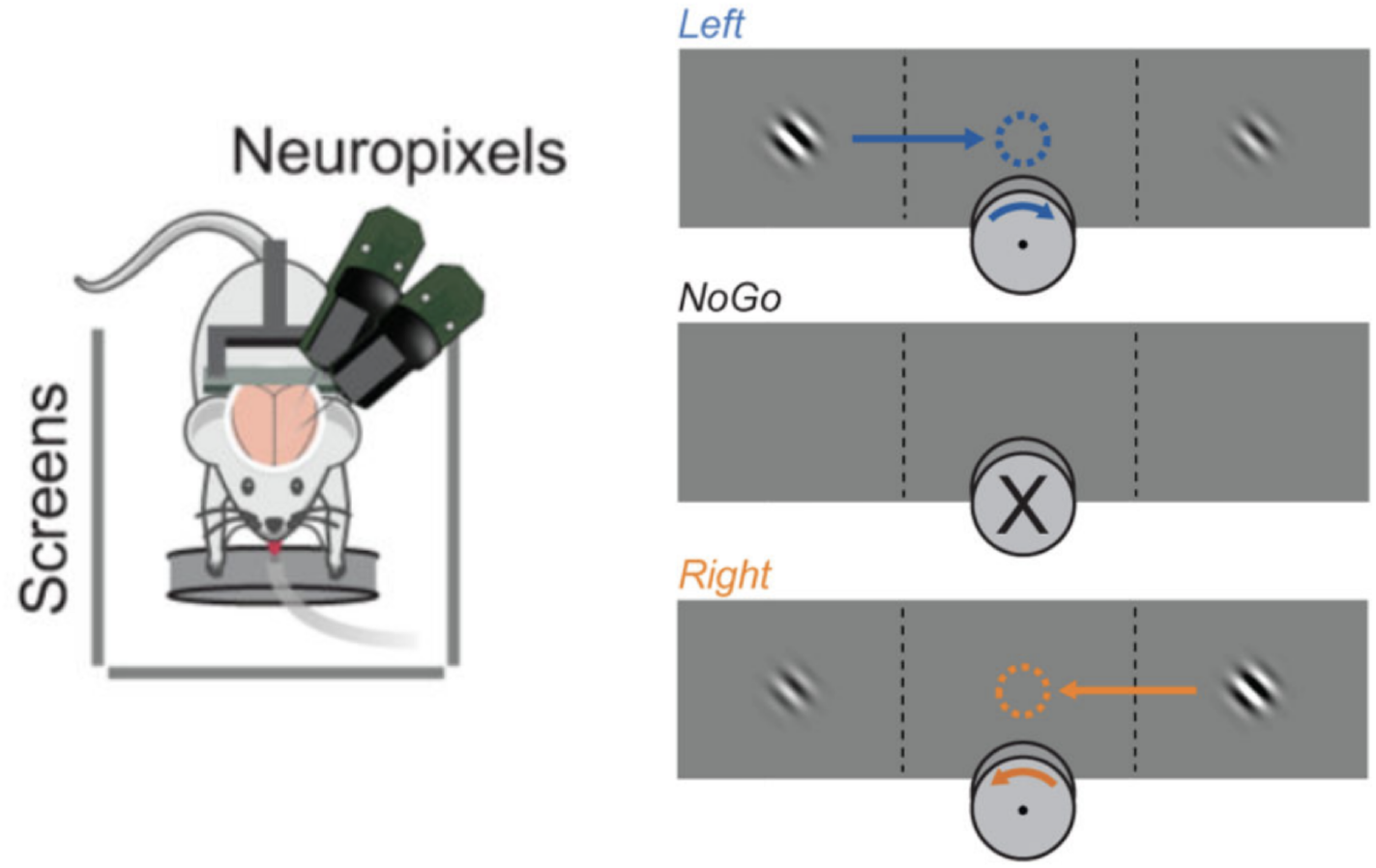
Experimental settings of dataset [20]

The experiment is done on multiple rats and repeated at different date. The dataset is sorted based on the name of the rat and the data of the experiment. Each dataset contains an array of values recording the neuron that spikes, the brain region it belongs to and the timing of the spiking event. We choose the portion of spiking events obtained from thalamus region.

### 2.3 Neuron model

The neurons in both sensory layers and population layers are modelled using the Hodgkin-Huxley model [21] with the equation

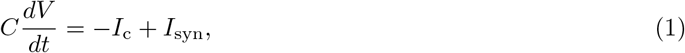

where *V* is the membrane potential, *C* = 1 *µ*F is the membrane capacity. The current is composed of channel current *I*_c_ and synaptic current *I*_syn_. The channel current is calculated as

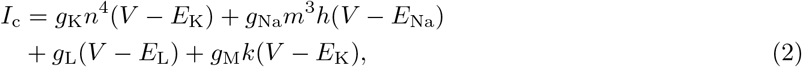

where *g* is the conductance with potassium (K and M), sodium (Na) and leaky (L) channel. The same notation is applied to the threshold voltage *E*_K_, *E*_Na_ and *E*_L_. The parameter values of conductance and threshold voltage are presented in Table 1 where + is used to denote excitatory neurons and *−* denotes inhibitory neurons. These values are designed to simulate hippocampal and cortical pyramidal neurons and inhibitory interneurons [22, 23]. The variables *m, n* and *h* are defined in the following form:

**Table 1.** Conductances and threshold voltages of ion channels (*E*_K_, *E*_Na_ and *E*_L_).

| Ion $i$ | $g_i^+$ (mS/cm<sup>2</sup>) | $E_i^+$ (mV) | $g_i^-$ (mS/cm<sup>2</sup>) | $E_i^-$ (mV) |
| --- | --- | --- | --- | --- |
| K | 5 | -100 | 9 | -65 |
| Na | 50 | 50 | 35 | 55 |
| L | $1 \times 10^{-2}$ | -70 | $1.33 \times 10^{-2}$ | -90 |
| M | $7.5 \times 10^{-2}$ | N/A | $9.8 \times 10^{-2}$ | N/A |

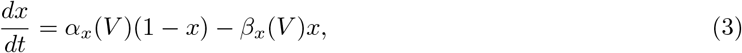

where *x ∈ {m, n, h}*. The function *α*_*x*_(*V*) and *β*_*x*_(*V*) for *m, n* and *h* is presented in Table 2, where 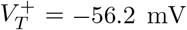 and 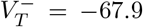 are the threshold voltages applied to each channel for excitatory and inhibitory neurons, respectively. The M-channel is a slow potassium channel with variable *k* calculated using a unique set of equation [24]:

**Table 2.**
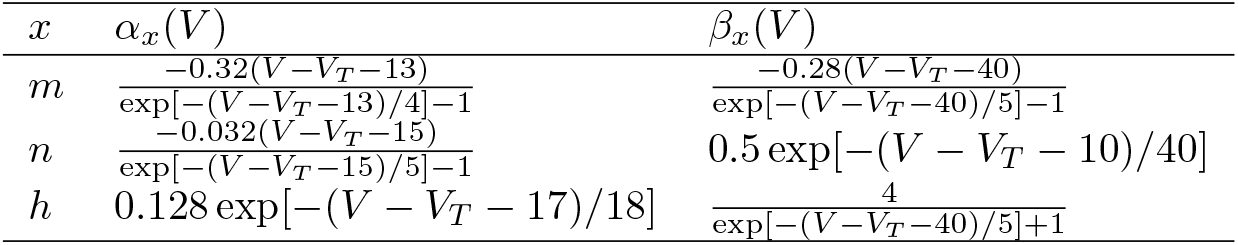
Functions of *α*_*x*_(*V*) and *β*_*x*_(*V*) for *m, n* and *h* in (1). *V* is membrane potential in mV.

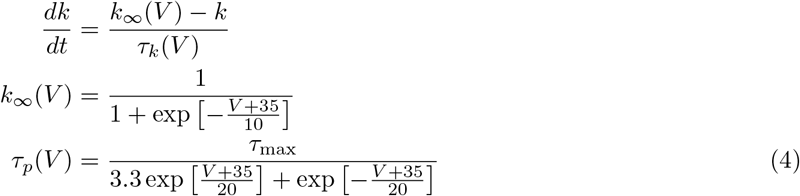

where *τ*_max_ = 4 s.

### 2.4 Synapse model

The synaptic current is further modelled as a combination of NMDA, AMPA and GABA synapses calculated with

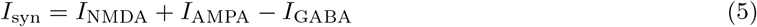

Each type of synaptic current is modelled according to Brunel and Wang [25]. The NMDA synaptic current of a neuron *q* is calculated as

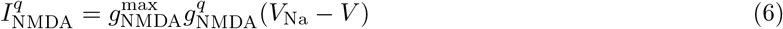

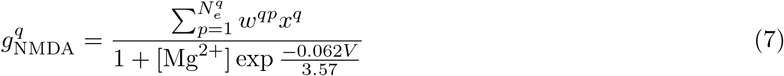

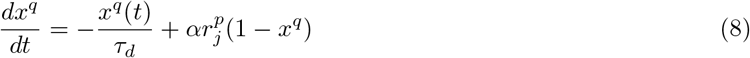

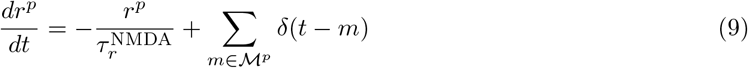

where 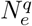 denotes total number of NMDA excitatory synapses; *ℳ*^*p*^ is a set of all spike timings. The Dirac delta summation 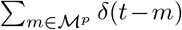 represents the spike trains. The value [Mg^2+^] represent the concentration parameter for magnesium ion. The weight connecting neuron *p* to neuron *q* is written as *w*^*qp*^. For AMPA and GABA synapses, the model takes on a similar form as

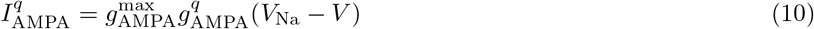

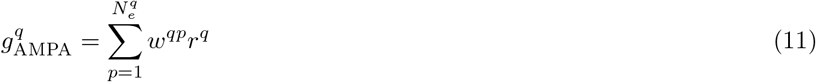

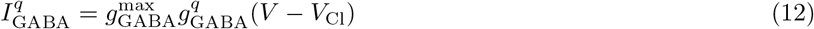

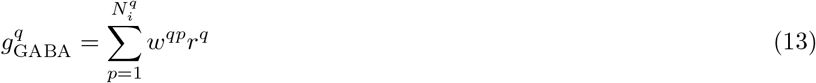

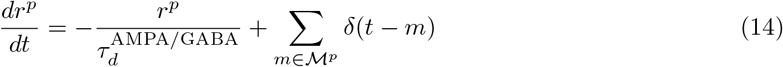

We also utilize the parameters given by Brunel and Wang [25], which are presented in Table 3 and Table 4.

**Table 3.** Maximum conductance and time constant used for synaptic modelling.

| Synapse $x$ | $g_x^{\max}$ (mS/cm<sup>2</sup>) | $\tau_d^x$ (ms) | $\tau_r^x$ |
| --- | --- | --- | --- |
| NMDA | 0.002 | 100 | 2 |
| AMPA | $7.5 \times 10^{-3}$ | 2 | N/A |
| GABA | $7.5 \times 10^{-3}$ | 10 | N/A |

**Table 4.**
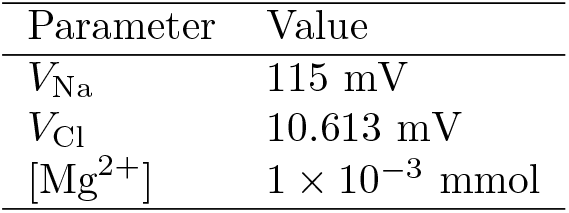
Threshold voltage and Mg ion concentration used for synaptic modelling.

| Parameter | Value |
| --- | --- |
| $V_{\text{Na}}$ | 115 mV |
| $V_{\text{Cl}}$ | 10.613 mV |
| $[\text{Mg}^{2+}]$ | $1 \times 10^{-3}$ mmol |

### 2.5 Spike-timing-dependent plasticity (STDP)

The weights *w*^*qp*^ from (9) and (14) are adjusted according to *spike-timing-dependent plasticity (STDP)* including both long-term potentiation (LTP) and long-term depression (LTD). Our simulation applies STDP equation proposed by Song et al. [26]:

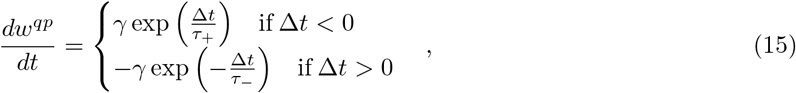

where the learning parameter *γ ∈* ℝ scales the weight adjustment. *p* is used to denote presynaptic neuron and *q* the postsynaptic neuron. We set *γ* = 10 and *τ*_+_ = *τ*_*−*_ = 20 ms. This definition allows our model to adhere more closely to biologically grounded equations.

We concentrate on synaptic weight changes within the population, fixing all other synapses to prevent complex neuronal network behaviours that may affect the resulting spike patterns. Therefore, STDP is restricted to intrapopulation synapses.

### 2.6 Simulation procedure

The model is implemented using BRIAN2 simulator [27]. The simulation runs for 1000 ms with step size of 0.5 ms. Each set of input data is provided to the model initialized in the same form. The initial membrane potential of all neurons is set to *−*60 mV. The synaptic weight is randomly initialized using a uniform distribution scaled by the number of inputs *N*. To test our proposal, we conduct the simulation under two conditions: one with learning enabled and one disabled. The detailed initialization function is outlined in Table 5.

**Table 5.** Synaptic weight initialization function using a uniform random variable *Y ~ U* (1, 1.1).

| *Presynaptic*→ | Input<sup>+</sup> | Input<sup>-</sup> | Sensory<sup>+</sup> | Sensory<sup>-</sup> | Population<sup>+</sup> | Population<sup>-</sup> |
| --- | --- | --- | --- | --- | --- | --- |
| Sensory<sup>+</sup> | $800Y/N$ | $800Y/N$ | N/A | N/A | N/A | N/A |
| Sensory<sup>-</sup> | $800Y/N$ | $800Y/N$ | N/A | N/A | N/A | N/A |
| Population<sup>+</sup> | N/A | N/A | $4Y/3N$ | $4Y/3N$ | $4Y/3N$ | $4Y/3N$ |
| Population<sup>-</sup> | N/A | N/A | $4Y/3N$ | $4Y/3N$ | $4Y/3N$ | $4Y/3N$ |
| *Postsynaptic*↑ |  |  |  |  |  |  |

The output spiking events of all neurons are plotted as raster plots in Figure 1b-d. Our focus is the population raster plot (Figure 1d). We observe over time that the population has varied spike timing. To quantify the desynchronization, we first separate “bands” of synchronization. The spike timings are grouped into local synchronous band (Figure 3) by the *z*-score outlier detection method [28]. The method calculates a so-called *z*-score using

**Fig. 3.**
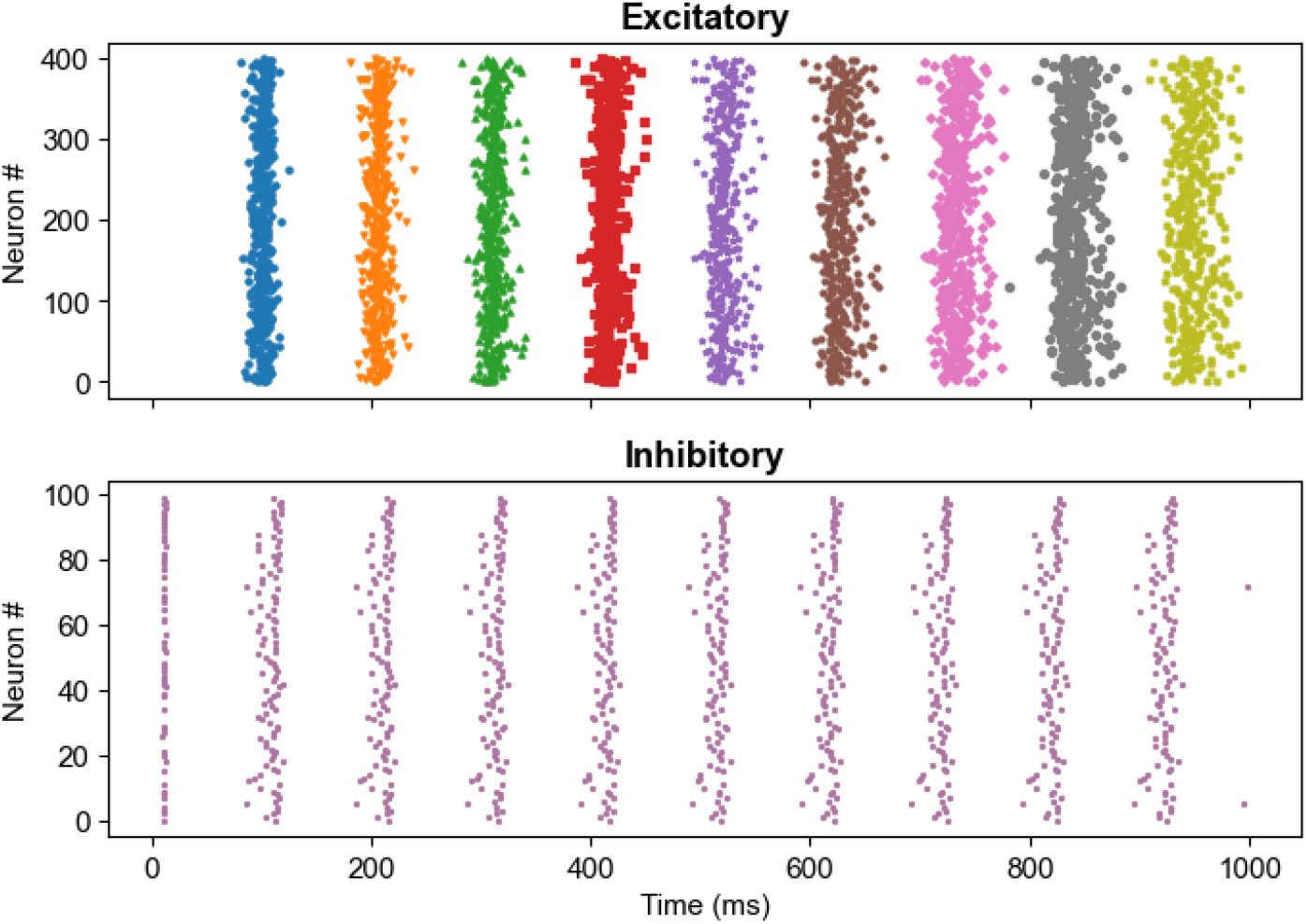
Synchronization bands (orange boxes) for grouping spike timings.

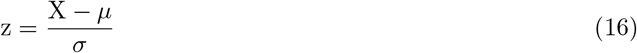

where *µ* is the average of the targeted points *X* and *σ* is the standard deviation. We find that when the score is above 7, the point can be fully separate into their respective synchronization band. The points of each band are then used calculate the mean and standard deviation of spike timing.

## 3 Results

### 3.1 STDP plays a critical role in desynchronization

Quantification of desynchronization is illustrated via pulse packet plots, that is, the plots of the standard deviation against the mean (Figure 4). The dotted lines in both plots outline the trend of spike timing variability. Colours of the lines indicate sessions of data input. The STDP-enabled network shows a tendency toward increasing variability as learning progresses (Figure 4a), suggesting the population undergoes a desynchronization phase whereas disabling learning under the same conditions yields mixture of both synchronization and desynchronization. However, slight resynchronization also occurs for some inputs shown by the bottom red line in Figure 4a. This resynchronization is fairly small on average roughly 4% of the preceeding variabiltiy. Nevertheless, under the influence of STDP, the spike timing variability at the end of the simulation is higher than that at the start of the simulation. The consistent observation of desynchronization during STDP learning suggests that STDP plays a critical role in desynchronization.

**Fig. 4.**
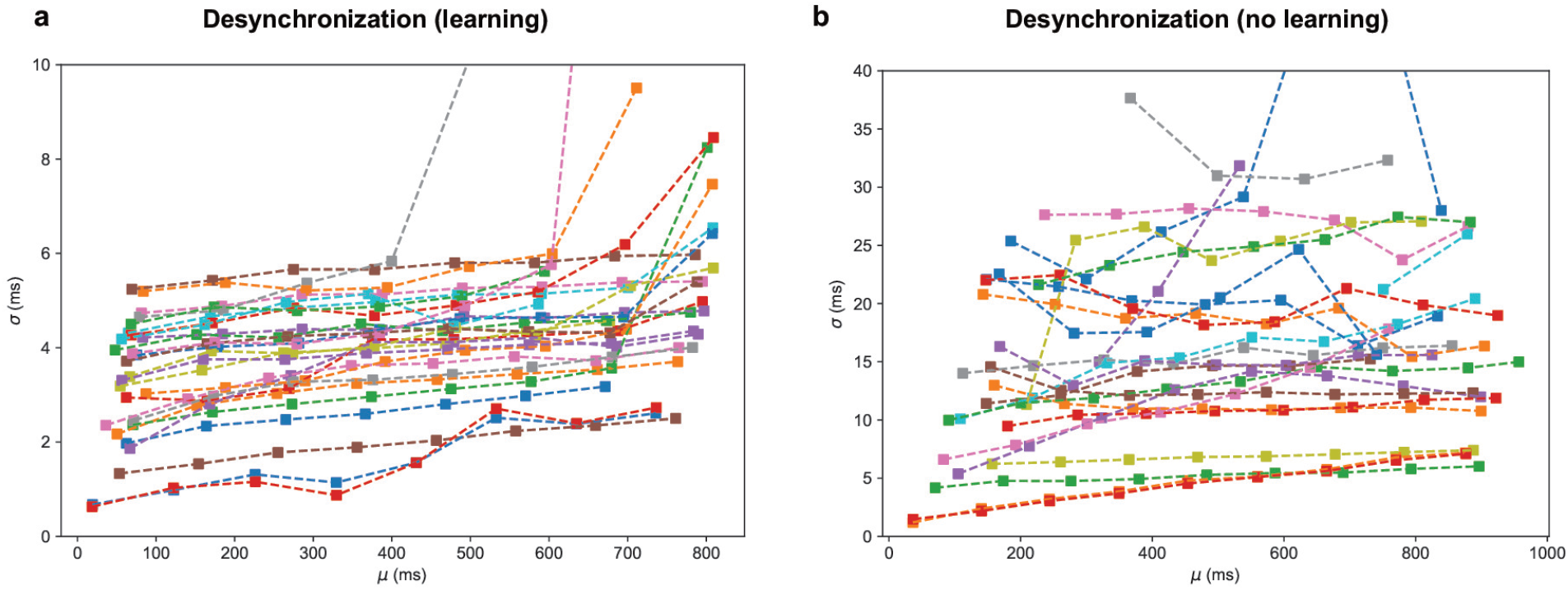
Desynchronization illustration: a: Pulse packet plot with STDP enabled. b: Pulse packet plot with learning disabled.

That STDP retains desynchronization has been pointed out via computational research [29, 30]. The interplay balances learning and desynchronization [31]. The balance is especially critical in terms of neurological symptoms such as epilepsy caused by overly synchronized neuronal behaviour and Alzheimer’s disease caused by overly desynchronized behaviour. The balancing is also explored in the realm of cognition and memory recalls [32]. We propose that cognition requires mutual balancing between STDP and learning. The proposal poses a question: Would desynchronization stablize at certain balancing points? Our previous result is insufficient to support the claim as the simulation time is too short to perceive substantial change. Therefore, we prolong the network simulation to inspect further the neuronal behaviour.

### 3.2 STDP and desynchronization are dynamically constrained

The network simulation is extended to 5 s with STDP enabled. The time is chosen so that the model can run sufficiently long with the limitation of computational memory. Since the dataset contains only the recordings lasting for 1 s, we augment the dataset by concatenating 5 recordings to form dataset inputs. The augmentation is applied to the dataset by concatenating the same recordings. In the experiment, they correpond to repeating the task with the same stimuli. We demonstrate the result using one session of recordings in Figure 5.

**Fig. 5.**
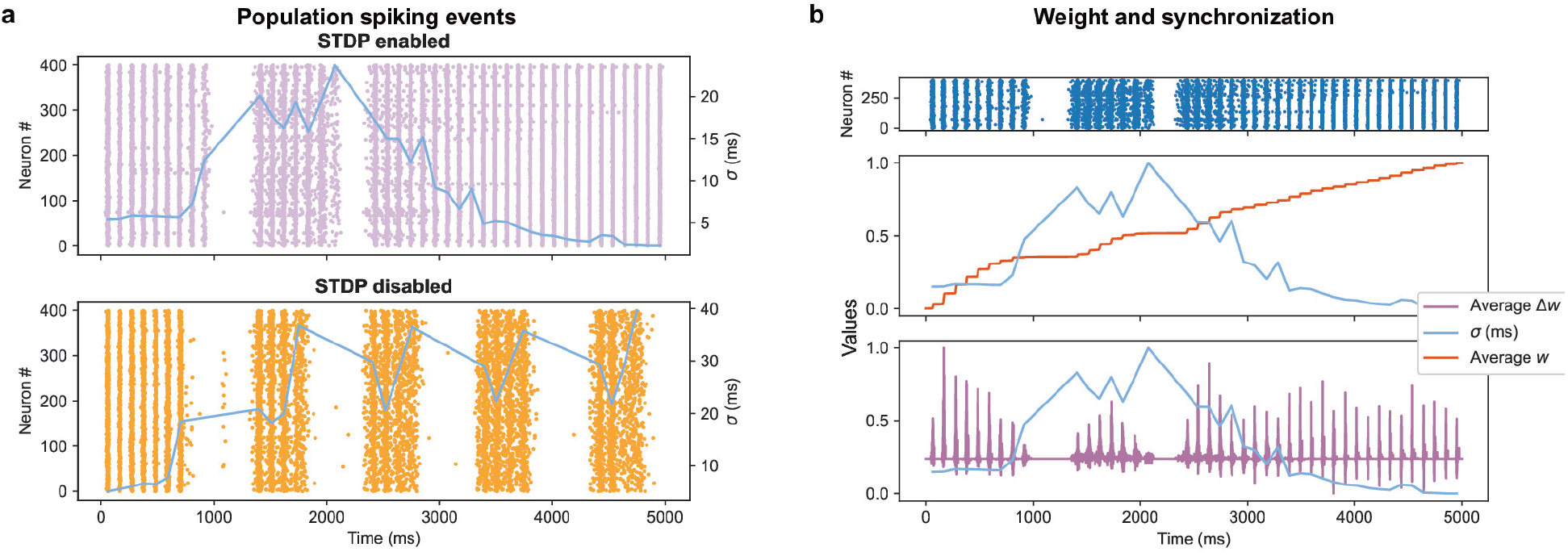
Example of simulation for 5s: a: Population neurons excitatory raster plot, with STDP enabled (top) and disabled (bottom). b: normalized pulse packet plot compared with excitatory raster plot (top), normalized weight *w* (middle) and normalized weight change Δ*w* (bottom).

The top plots in Figure 5a outline the spiking events with STDP enabled and disabled. While the plot contains synchronized spiking events, it also periodically introduces a synchronized nonspiking time period. Further quantification of the synchrony is done via normalized pulse packets (blue lines in the middle and bottom plots of Figure 5a and b). In Figure 5a, comparison between the top and bottom plot reveals that without STDP, desynchronization events grows much higher with STDP enabled, suggesting that STDP is essential in regulating desynchronization and maintaining the level of synchrony. In Figure 5b, we align all data using normalization in the plot for trend comparisons. Besides the previous increased desynchronization for the first second, we observe synchrony change in longer periods. To our surprise, the level of synchrony does not fluctuate around a fixed point but rather rapidly increases (synchronized) and decreases (desynchornized). The global trend of this behaviour is a fluctuation that drives toward synchronization eventually. It is worth noting that the beginning of such fluctuation is preceeded by the first gap period. In the bottom plot from Figure 5b, the desynchronization is no longer observed with the disappearance of the gap period.

The rollercoaster-shaped change of synchrony is accompanied by STDP weight adjustments. Here, we plot the average weight (orange lines) as well as the average weight change (purple lines) of all available synapses in Figure 5b. The STDP intensity is limited by the level of desynchronization. The effect is especially evident in the bottom plot of Figure 5b from 2 s to 5 s. In combination with (15), decreased synchronization limits the overall adjustment intensity imposed by STDP. While initial adjustment (up to 1 s) of STDP tending toward neuronal desynchronization, the later adjustment of STDP generates a more diverse neuronal behaviour. Within continued STDP adjustments (such as 1 s to 2 s of bottom plot Figure 5b) include both desynchronization and resynchronization. What is more interesting is that compared to the desynchronization phase the resynchronization begins right after the longer gap where neurons are sparsely spiking. The ensu synchronous spiking events transition from resynchroning to desynchronization.

Contrary to our expectation, the neuronal variability does not converge toward a fixed stable point but rather fluctuate during STDP adjustment. However, this does not counter the mutual feedback between STDP and desynchronization. In fact, the switch from resynchronization to desynchronization is evidence suggesting such feedback exists during active STDP learning. Nevertheless, we are puzzled by the longer synchornization period starting after 3 s. The behaviour seemingly suggests that the population enters a regular firing mode. Different datasets reveal that the timing of the entry varied greatly (from 2 s to 4 s). It is yet to be understood what causes such synchronization.

### 3.3 Summary

In summary, we propose that STDP plays a critical role in neurons desynchronization. We start by confirming that early stages to learning creates a desynchronization effect within the population. This outcome leads to the proposed feedback control between STDP and synchronization that eventually could stablize at a reasonable variability. To rationalize this proposal, we extended the simulation time to 5 s. The result verifies that desynchronization could reduce the overall intensity of STDP. However, we do not observe variability converges toward a fixed point. Rather, inclusion of STDP supresses the desynchronization by switching population between desynchronization and resynchronization.

## 4 Discussions

We have demonstrated via computational means that STDP and desynchronization mutually affect one another. The interaction between STDP and desynchronization are thus summarized in two arguments:

- STDP tunes population by switching from synchronization to desynchronization.
- Reduction in synchronization of spikes lowers the intensity of STDP (in *w*^*qp*^).

The combination of these two arguments form an adjustment loop that limits each other. The adjustment loop does not tune the synchronization toward a stablizing value. Rather, it drives a repetitive fluctuation contained within a range.

As important as synchronization, desynchronization plays a crucial role in memory encoding and retrieval [7]. This statement is supported experimentally and computationally [8, 33]. Hippocampal synchronization and neocortical desynchronization underlie the episodic memory formation [34]. Studies in recent years have demonstrated desynchronization brain state is related to learning and can be used to study memory acquisition [35]. Our computational study agrees with these previous results that learning is connected to desynchronization and can be tied to memory encoding and acquisition. Under the same stimulus response, the fluctuation signifies a dynamical encoding scheme of neurons. Our simulation mimics hippocampal response within the brain. The synchronized activity within the region is generally viewed as the encoded information. An example of such encoding is the “place” cell, a type of neuron that activates to indicate positions [36]. Our result shows that STDP creates a dynamical synchronization, indicating that the learning process allows variability in encoding information. Theory of neural self-information coding has pointed out through biological studies that population variability is large during cognition [37], yielding validity to the changing synchrony during learning period.

Beyond the consistency of our results with prior findings, our theory also offers an explanation for the longstanding synchronization–desynchronization conundrum. Studies of neural synchronization emphasize learning arising from coordinated neuronal activity, whereas information theory highlights the role of desynchronization [2]. The example of place cells where neuron activation encodes the spatial location, suggests that neurons require varied signals for varied positions. This reasoning supports the information-theory side of the conundrum. However, knowledge should be acquired via STDP during synchronization. In terms of place cells, neurons need to synchronize together to start the learning process yet desynchornizes to allow flexibility of representation. Our results provide an alternative perspective by demonstrating that learning can implicitly promote synchronization and desynchronization. Specifically, our computational model shows that STDP-driven synaptic weight adjustments initially desynchronizes the population, which in turn attenuates the strength of further learning. At higher level of desynchronization, STDP is also capable of switching between synchronization and desynchronization. In this framework, learning governs both synchronization and desynchronization. Combining with the desynchronization that feeds back to regulate synaptic plasticity, this reciprocal interaction offers a coherent resolution to the synchronization–desynchronization conundrum.

Based on this process, we propose that STDP learning behaviour interacts with neurons in the following way. At the onset of an input, an encoded representation initiates a group of synapses to perform learning. The spike rate and timing start to align for selected neurons. As learning grows active, neuron spiking events gradually desynchronizes and thus learning reduces the intensity. Concurrently, the spike rates of the targeted neurons start to diversify. As the diversification reaches certain level, STDP starts to tune the level of synchrony to counteract the over-desynchronization and over-synchornization and slowly returns learning back to a synchronized state. These results establish a link between STDP and information encoding with the help of synchronization and thus supporting the existence of a control loop coupling synchronization, desynchronization, and synaptic plasticity.

